# A comprehensive automated pipeline for human microbiome sampling, 16S rRNA gene sequencing and bioinformatics processing

**DOI:** 10.1101/286526

**Authors:** LW Hugerth, M Seifert, AAL Pennhag, J Du, MC Hamsten, I Schuppe-Koistinen, L Engstrand

## Abstract

The advent of affordable high-throughput DNA sequencing has opened up a golden age of studies in the human microbiome. In order to understand the role of the human microbiota, standardized methods for large-scale, population-level studies are needed to avoid underpowered or poorly designed studies. The biggest bottlenecks to population-level microbiomics are sample collection, storage and DNA extraction. Here, we describe a flexible automated approach to process intestinal biopsies, fecal samples and vaginal swabs from sample collection to OTU table. We have evaluated storage conditions, DNA extraction methods, PCR strategies and bioinformatic pipelines for these three sample types, and present here a set of guidelines and best practices for each of these steps.

## INTRODUCTION

The advent of affordable high-throughput DNA sequencing has opened up a golden age for studies in the human microbiome. Sampling strategies covering hundreds of subjects [1–3] or comprehensive spatial or temporal sampling of a few individuals are now possible [4, 5]. The explosion of studies in microbiomics combined with the rapid adoption of this research field by researchers of various backgrounds has increased the risk of publishing underpowered or otherwise ill-designed studies. Today, large-scale, population or hospital-based studies are often needed to increase our understanding of the role of the microbiome in various diseases. With the time from sample preparation to sequencing results now counted in days, the biggest bottleneck to population-level microbiomics are now sample collection, storage and DNA extraction. In addition to being expensive and time-consuming, a sub-optimal DNA extraction can lead to severe biases in the study, and ultimately false conclusions [6].

One of the most common source materials for human microbiome studies are faecal samples. The large intestine has the greatest concentration of bacteria in the human body [7] and the fecal microbiome has been linked to a wide variety of gastrointestinal [8, 9], metabolic [10, 11] and even neurological conditions [12, 13]. Faecal samples can also be collected at a moderate cost and non-invasively, making this a suitable and popular target for studies of the human microbiome.

One problem with faecal samples is that they represent a large bulk volume which is not in direct contact with the host’s mucosal lining. While it is reasonable to assume that products of microbial metabolism in the luminal space, such as short chain fatty acids, can affect host physiology [14] it is also true that bacteria living in intimate association with the mucus layer in the gut lining likely have a stronger effect in modulating the host’s immune response [15]. As these niches present quite different selection pressures, bacteria found attached to the gut lining form a clearly separate community from those in the luminal space [16] and can only be queried through the use of gut biopsies.

While biopsies obtained from the gastrointestinal tract allow the investigation of bacteria in tighter attachment and deeper layers than a simple swab, they are a different type of material compared to faecal samples. Firstly, contrary to a faecal sample, the vast majority of the DNA in a typical biopsy is from the host, rather than bacterial. Furthermore, the bacteria are living in a complex three-dimensional biofilm, which might be harder to disrupt. Available commercial kits for selectively removing human DNA are inappropriate when the study design includes host genotype or eukaryotic microbe profiling and might inadvertently remove part of the microbial diversity. Consequently, the risk of introducing bias is obvious. Therefore, a DNA library preparation method of broad applicability needs to be robust to overwhelming proportions of host DNA.

Other host surfaces, while not particularly rich in host DNA, present a chemically complex extracellular matrix, which can hinder DNA purification, and, in some cases, a relatively low bacterial cell count. Sputum and saliva are good examples of this, as is vaginal mucus. The latter is a particularly important target in gynecological screening [17] and might be an important prognostic tool for obstetric and neonatal health [3]. A human microbiome pipeline of general applicability should also apply to mucus-associated microbes.

Faecal and mucus samples are relatively easy to retrieve and can often be collected by the research subject at home. This raises the issue of the correct storage procedure for these materials. Left at room temperature, a bacterial community can present significant shifts after only a few minutes, due to overgrowth of oxygen-tolerant microorganisms. Therefore, it is crucial to assess bacteriostatic and preservation strategies. It is important to consider cost, ease of use for the research subject, non-toxicity, quality of sample preservation and compatibility with downstream applications.

Once a good procedure for sample collection and DNA extraction has been established, the next challenge for an amplicon-based study (eg 16S rRNA gene surveys) is an appropriate PCR strategy [6]. The most crucial choice is the selection of broad-taxonomic range primers compatible with the target community [18]. The thermodynamic characteristics of the primer pair will compound to the biases, through preferential annealing or incomplete melting of GC-rich sequences [19]. It is also crucial to work under appropriate molecular biological conditions, considering that a single molecule of contaminating DNA can be amplified to 1000 copies after only ten PCR cycles. Reducing the number of PCR cycles can thus ensure a less biased picture of the community. Avoiding intermediate cleaning steps also reduces the risk of sample spillover and cross-contamination. Finally, before sequencing, sample pooling is another sensitive step, where the depth of sequencing for each sample is determined.

Challenges still remain after DNA sequencing, though, since bioinformatic processing presents its own set of challenges [6]. In the early days of metabarcoding, clustering was necessary, partly to collapse erroneous sequences to true biological diversity and partly to make sequence clusters (operational taxonomic units, OTU) large enough for quantitative statistical methods to apply. The latter is not an issue with current high-throughput technologies, which typically provide sufficient data for much finer clustering. Sequencing errors and minor biological variation, however, do artificially inflate the number of unique OTU, compared to the true number of sequences or strains [20]. However, many modern error correction strategies eschew the need for an *a priori* similarity cut-off [21–23]. This is crucial for vaginal microbiome studies. While most vaginal communities are dominated by Lactobacilli, it has been shown that communities characterized by a dominance of *L. iners* are less stable than those dominated by *L. crispatus* or *L. gasseri*. A species-level identification is therefore crucial. This issue is even more extreme for skin microbiome studies, where it is necessary to differentiate *Staphylococcus epidermidis* from *S. aureus*, although they differ by only 14 bp difference over their whole 16S rRNA gene, and by only 2 bp in the commonly used 341-806 region.

Using a high-resolution error correction method requires a taxonomic assignment strategy of compatible sensitivity. While there are good tools available for general taxonomic surveys [24, 25], specific research questions sometimes require custom-made taxonomic approaches [26, 27].

Here, we describe a flexible automated approach to process intestinal biopsies, faecal samples, vaginal swabs and saliva samples, from collection to OTU table. We have eavlueated sample storage, DNA extraction, PCR and bioinformatic pipelines for these three sample types. We present a set of guidelines and best practices for each of these steps, which was shown to also work for saliva samples and can likely be extended to other swabs and bodily fluids.

## RESULTS AND DISCUSSION

### Sample collection and storage

Biopsies are necessarily taken in a hospital or clinic, and therefore present the best conditions for sample preservation. Fresh biopsies are sometimes placed in filter paper after extraction. We notice that, in this case, the paper should also be submitted to DNA extraction, together with the tissue. The same thing applies to biopsies or swabs preserved in liquid medium, where both the liquid and the solid fractions should be taken for extraction.

We have compared freezing fresh samples at −80°C or freezing them in three distinct preservation media, RNALater, Allprotect and DNA/RNA Shield (see Methods for details). RNALater did not give as high DNA yields as the other two methods (data not shown). DNA/RNA shield is compatible with all downstream steps and presented excellent storage characteristics, as described below, and was therefore selected for further sampling.

Faecal samples are often collected at the patient’s home, since it can be difficult to produce the material at the time of clinical examination. This means that a −80°C freezer is not available, although a −20°C often is. Even then, there is risk of thawing during transportation, so a preservation medium might be required. We compared two faecal samples from the same healthy volunteer: one sample was immediately frozen at −20°C, while the other was divided into two fractions, whereof one was placed in DNA/RNA shield and the other kept dry at −20°C, simulating patient self-collection. The next day, the sample was briefly thawed and homogenized in DNA/RNA shield. Then, one fraction of the homogenate was immediately extracted while the others were kept for 8 days at −20°C or −80°C. There was a clear difference in the diversity of the sample kept in DNA/RNA shield as compared to the one frozen dry **(fig 1a).** We hypothesize that this is due to overgrowth of aerotolerant microbes prior to freezing and possibly during the intermediate thawing, leading to a skewed community. The same did not happen in DNA/RNA shield, which inactivates bacteria in seconds to minutes. Further preservation of the sample for up to 8 days in either −20°C or −80°C didn’t affect the inferred community, showing that getting the sample quickly from the patient’s home to the clinic is of minor concern, as long as the sample is efficiently inactivated **(fig 1b)**.

**Figure 1.**
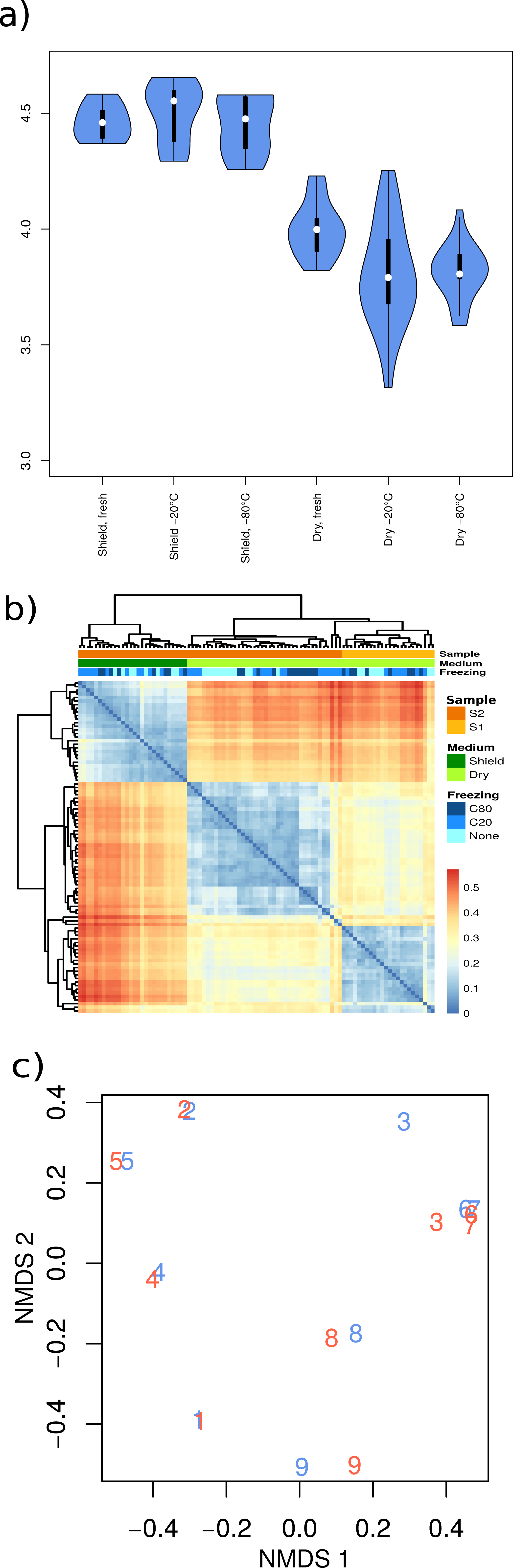
Effects of sample collection and preservation. (a) Fecal samples frozen without a preservation media display reduced Shannon entropy. Shield: sample preserved immediately in DNA/RNA-shield. Dry: sample kept without a preserving medium. Fresh: sample kept overnight at −20°C prior to extraction. −20°C, −80°C: samples kept in each of these temperatures for 8 days, prior to extraction. (b) Preserving samples with or without a preservative has a larger impact in their microbial composition than sampling in separate occasions. Clustering based on Bray-Curtis divergence. S1, S2: sampling occasions. Shield, dry: sample preserve with or without DNA/RNA-shield. C80, C20, None: samples kept for 8 days at −80°C, −20°C or extracted as soon as possible. (c) Samples taken by the patient (blue) or by a professional (red) cluster together, indicating that sample self-collection is a viable research approach. Subject 3 had a Lactobacilli-dominated vaginal community, but in the self-taken sample a 1% presence of other Gram-positive bacteria drives the shift away from its replicate. NMDS on 2 dimensions based on Bray-Curtis distances.

Vaginal swabs can also be collected at home. We observed no difference between samples collected by the patients themselves or samples collected by midwives in a clinical setting **(fig 1c)**. Swabs were kept at room temperature in DNA/RNA shield for up to three days before being frozen at −80°C. Since bacteria might remain attached to the swab or come loose in the medium, both of these were included in the DNA extraction.

### DNA extraction

A good DNA extraction strategy should be effective, have minimal hands-on time, be non-toxic, automatable and highly reproducible. Two commercial solutions were assessed, MoBio’s PowerMag and Zymo Research’s ZR-96 Genomic DNA MagPrep. No conclusive difference in data quality was found between the MoBio and the Zymo approaches. However, since the latter doesn’t require centrifugation, making it more suitable for automation, it was selected for further optimization. The MoBio kit also uses β-mercaptoethanol, a strong smelling solvent which may require the use of a chemical hood.

Bead-beating is a crucial step for homogenizing samples, destroying extracellular matrix and opening up cells with tough walls, such as Gram-positive bacteria. We therefore paid special attention to this issue, considering the size of the beads used, the duration of bead-beating and the amount of starting material. All samples were homogenised in an initial bead-beating procedure. After digestion with lysozyme and proteinase K, we assessed whether an extra bead beating step, with finer beads, could yield increased recovery of Gram-positive bacteria. We found that the extra bead-beating increases slightly the DNA yield for all samples, but doesn’t make a large difference in their overall composition (**fig 2).** Due to considerations on time, cost and contamination risk, this additional bead-beating step was not performed in subsequent experiments.

**Figure 2.**
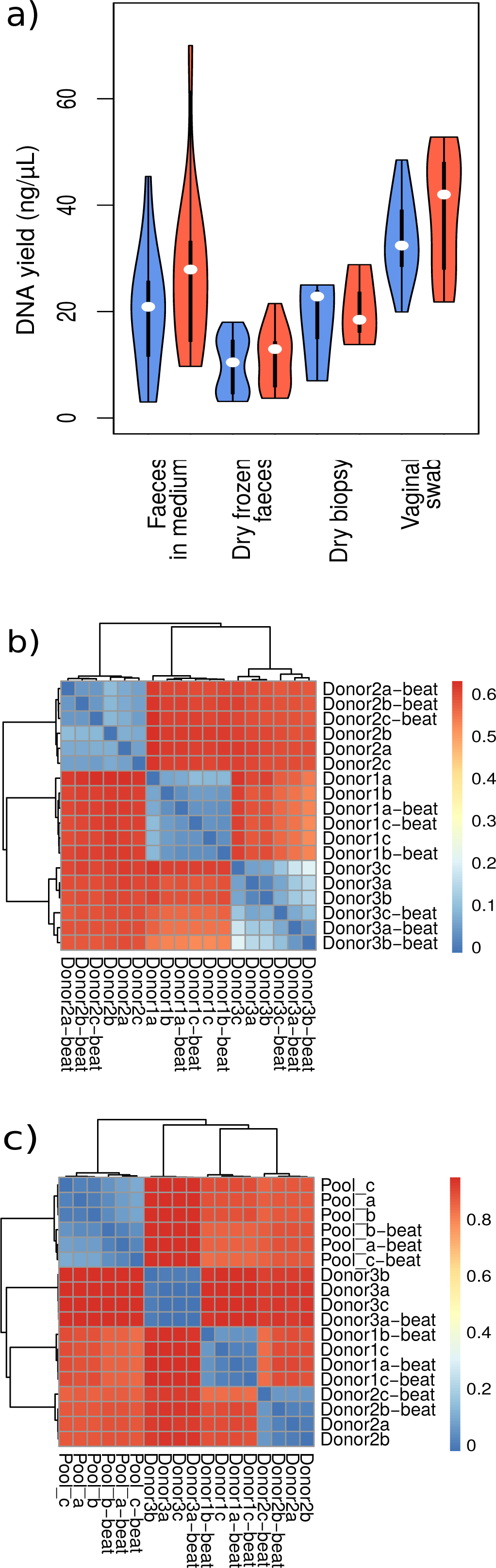
(a) Violin plot depicting the DNA yield of different sample types with and without the additional bead beating step. Orange violins have the additional bead-beating step, blue ones do not. (b and c) Heatmaps displaying Bray-Curtis divergence between samples treated with an additional bead-beating step or not; “beat” marks the samples submitted to this treatment. Donors are numbered arbitrarily in each panel and have no correspondence across them. The pool comprises several donors, not only the individuals otherwise included in the figure. (b) Fecal samples (c) Vaginal swabs.

In addition to the physical steps of heating and bead-beating, a chemical digestion of bacterial cell walls is needed for DNA extraction. Besides the proteinase K step, we have assessed the efficiency of pure lysozyme compared to Molzym’s BugLysis kit. No difference was observed, and Lysozyme was selected for further optimization. The time and temperature of incubation in lysozyme was then optimized, showing the reaction to be fairly temperature insensitive to temperature and time, with an incubation of 30-60 minutes functioning equally well (**fig 3)**, after which the quality of the extracted DNA might fall.

**Figure 3.**
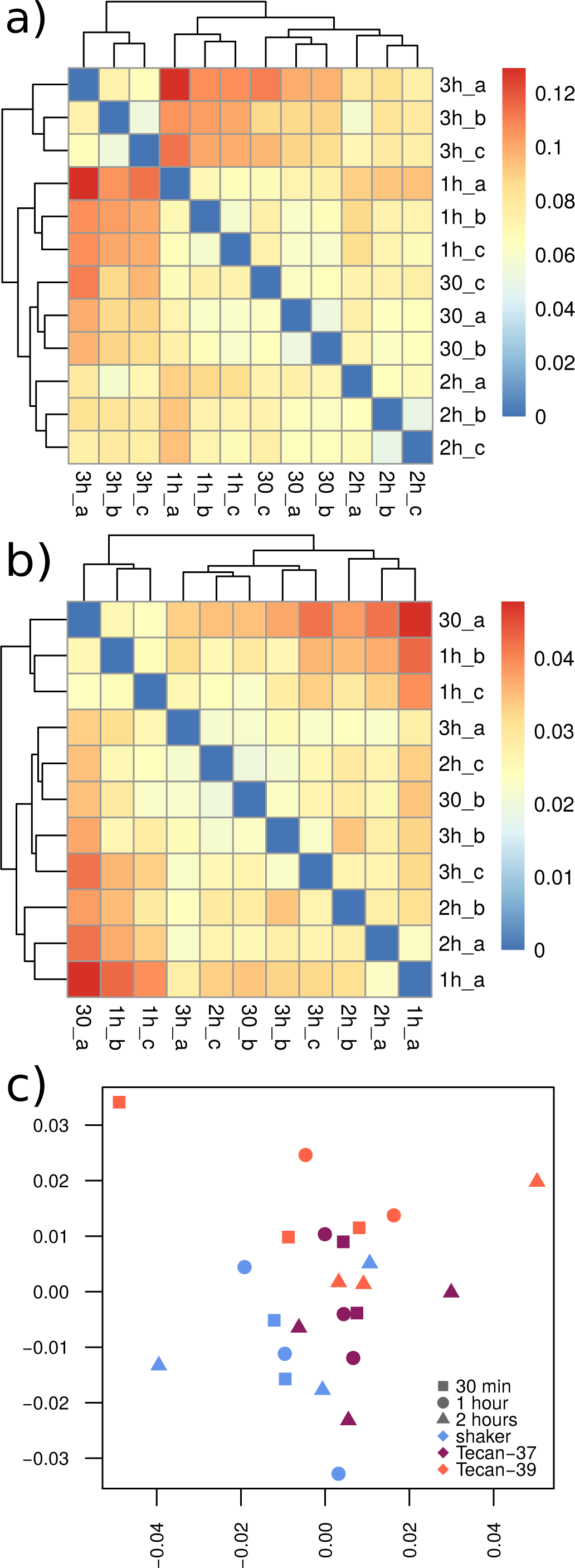
(a and b) Heatmaps displaying Bray-Curtis divergence between (a) fecal samples and (b) vaginal swabs incubated in lysozyme for 30, 60, 90, 120 or 180 minutes. (c) NMDS of vaginal samples that were heated at 37°C or 39°C in the TECAN robot or at 37°C in a separate shaker. While all figures show some clustering per technical triplicate, the distance between samples is minimal and the effect is not linear with time.

The final step of DNA extraction is to release the pure DNA molecules into solution for storage and downstream applications. For this step, three possibilities were considered: milliQ water, Tris-Cl/EDTA (TE) and Tris-Cl (EB). Since water doesn’t preserve DNA quality as well as the buffers, and EDTA is incompatible with certain molecular applications, such as the use of restriction enzymes, EB was selected as the elution buffer.

Finally, for shotgun metagenomics or eukaryotic marker gene amplification, it is tempting to deplete the sample from host DNA, specially from biopsies. However, we have found that the Molzym treatment for human DNA removal, developed for blood samples, also removes bacterial DNA and shows preferential removal of specific clades, most notably Clostridiales, when applied to intestinal biopsies (**fig. 4**).

**Figure 4.**
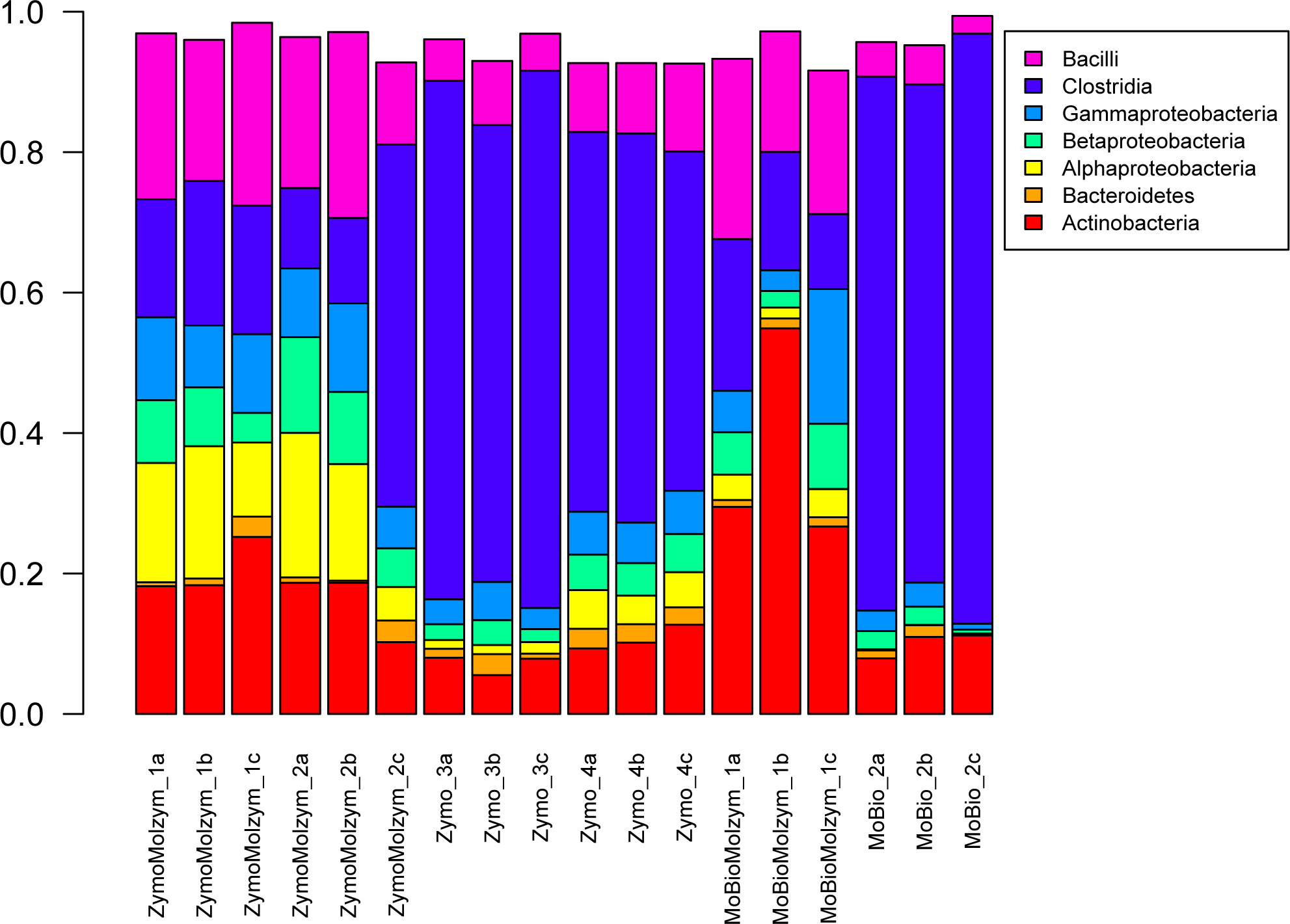
Removing human DNA selectively removes DNA from certain clades. Samples are extracted with ZymoBiomics or MoBio kits as stated, and treated with Molzym human DNA removal as stated. The Molzym treated fecal samples show decreased prevalence of Clostridia.

During the preparation of this manuscript, Zymo Research phased out the ZR-96 Genomic DNA MagPrep kit and replaced with the kit Quick-DNA MagBead Plus. The DNA yield for this kit is generally higher, specially for vaginal samples, but does not present a higher level of background DNA (**suppl. fig. 1a)**. This difference in DNA yield is likely due to a better rupture of Gram-positive cells, such as Lactobacilli.

The alpha-diversity for the same biological samples extracted with either kit is highly similar, and the effect on beta-diversity is negligible (**suppl. fig 1b-c**). While we do not recommend mixing extraction protocols during the same study, the data presented in this work holds true regardless of which of these two kits is used.

### 1-step PCR amplification

A common approach to amplicon library preparation is to run two PCR reactions, whereby the first one targets the region of interest and the second one adds identifying barcodes and any other sequences necessary for the desired sequencing platform. However, this approach is very vulnerable to contamination and errors. We have therefore developed a 1-step PCR procedure that eliminates the need for an intermediate cleaning step, which is described in detail in the Methods’ section.

Although most samples maintained coherence between the 1-step and 2-step procedure **(fig 5a)**, for some samples the 2-step PCR sample has greatly decreased diversity (**fig 5b)**, suggesting over-amplification as a consequence of the additional number of PCR cycles. Furthermore, the 1-step procedure decreased the number of sequences with low-resolution taxonomic assignment, suggesting a decrease in chimeric, erroneous and non-bacterial contaminating sequences (**fig 5c-d)**. By eliminating a cleaning step, this procedure also reduces the per-reaction cost significantly.

**Figure 5.**
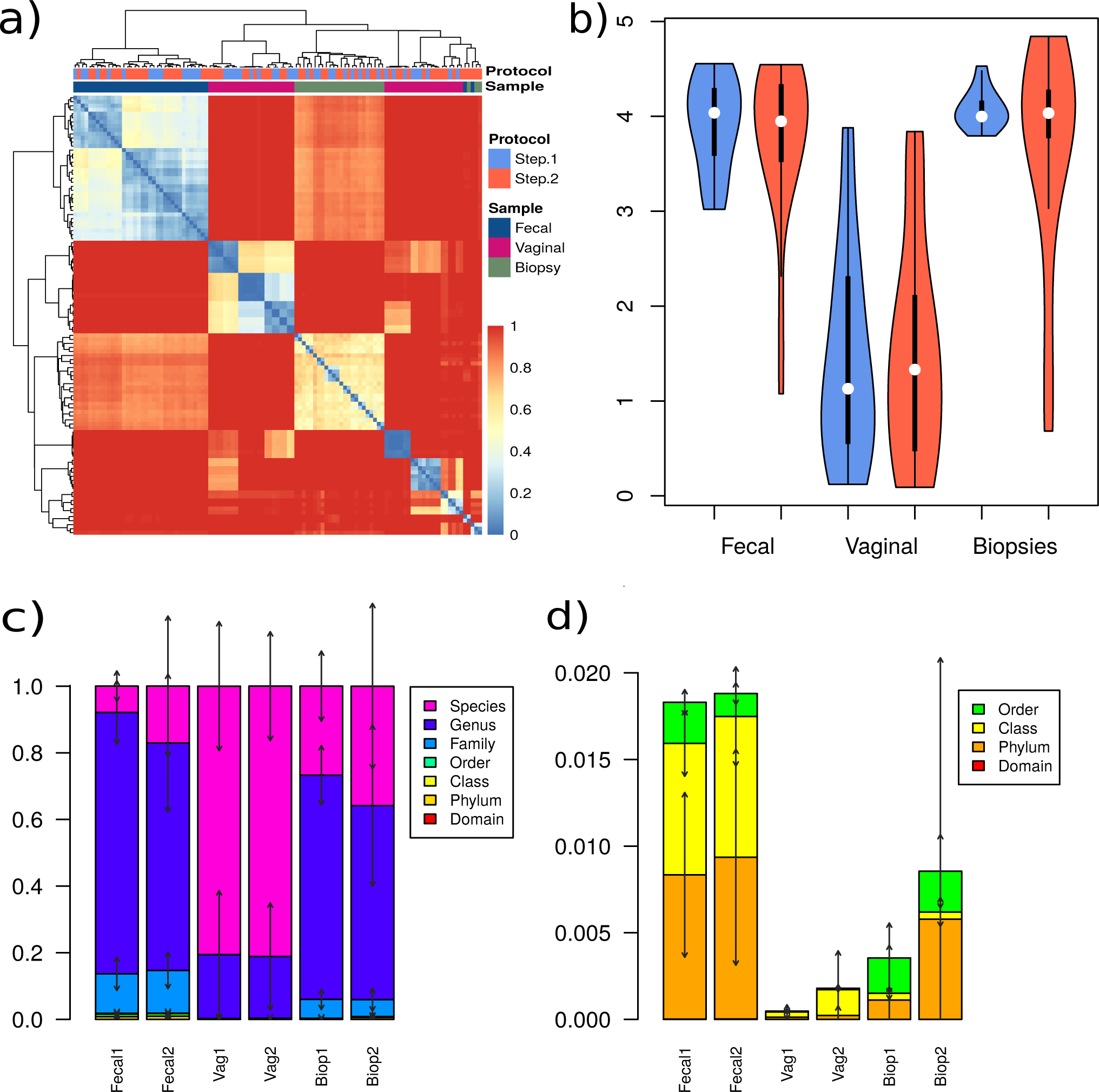
1-step and 2-step PCR procedures yield mostly comparable results, but 2-step shows signs of over-amplification, consistent with the larger number of PCR steps used. (a) Heatmap of Bray-Curtis distances between vaginal, faecal and gut biopsy samples amplified with a 1-step or 2-step PCR procedure. Samples cluster firstly according to sample origin and then, mostly, according to individual sample, with 1-step and 2-step clustering close to each other. Vaginal samples are split between extreme Lactobacilli-dominated samples and more diverse communities. The outgroup of 2-step procedures on the right edge of the graph corresponds to samples with reduced diversity and possible over-amplification (b) The alpha-diversity of most samples is preserved across 1-step and 2-step procedures, but on a few occasions biopsy and faecal samples are dominated by a single clade on the 2-step procedure, likely due to over-amplification. (c) The depth of taxonomic assignment is similar for 1-step and 2-step procedure, with a tendency for more reads with species-level assignment in the 2-step procedure. This is due to a lower proportion of rare, poorly classified OTU. (d) A zoom-in into figure (c), focusing on higher-order taxonomic levels. In this case, there’s a tendency of increase in the proportion of reads that cannot be classified more deeply than the phylum and class levels with the 2-step procedure, possibly due to the accumulation of PCR-errors and possibly chimeras.

### Depth of sequencing

Given a limited budget, sequencing depth is always a trade-off between deeply understanding a few samples for full biological insight of them versus having enough replicates and controls to reach statistical significance. A standard approach for assessing sufficiency of sequencing depth is through rarefaction curves, which answer the question of whether every unique gene tag in the target population has been seen at least once. Rarefaction curves for intestinal biopsies, fecal samples, vaginal swabs and saliva samples show that at 50,000 reads, the sequencing effort is not enough to uncover the full richness of the samples studied **(fig 6a-d)**.

**Figure 6.**
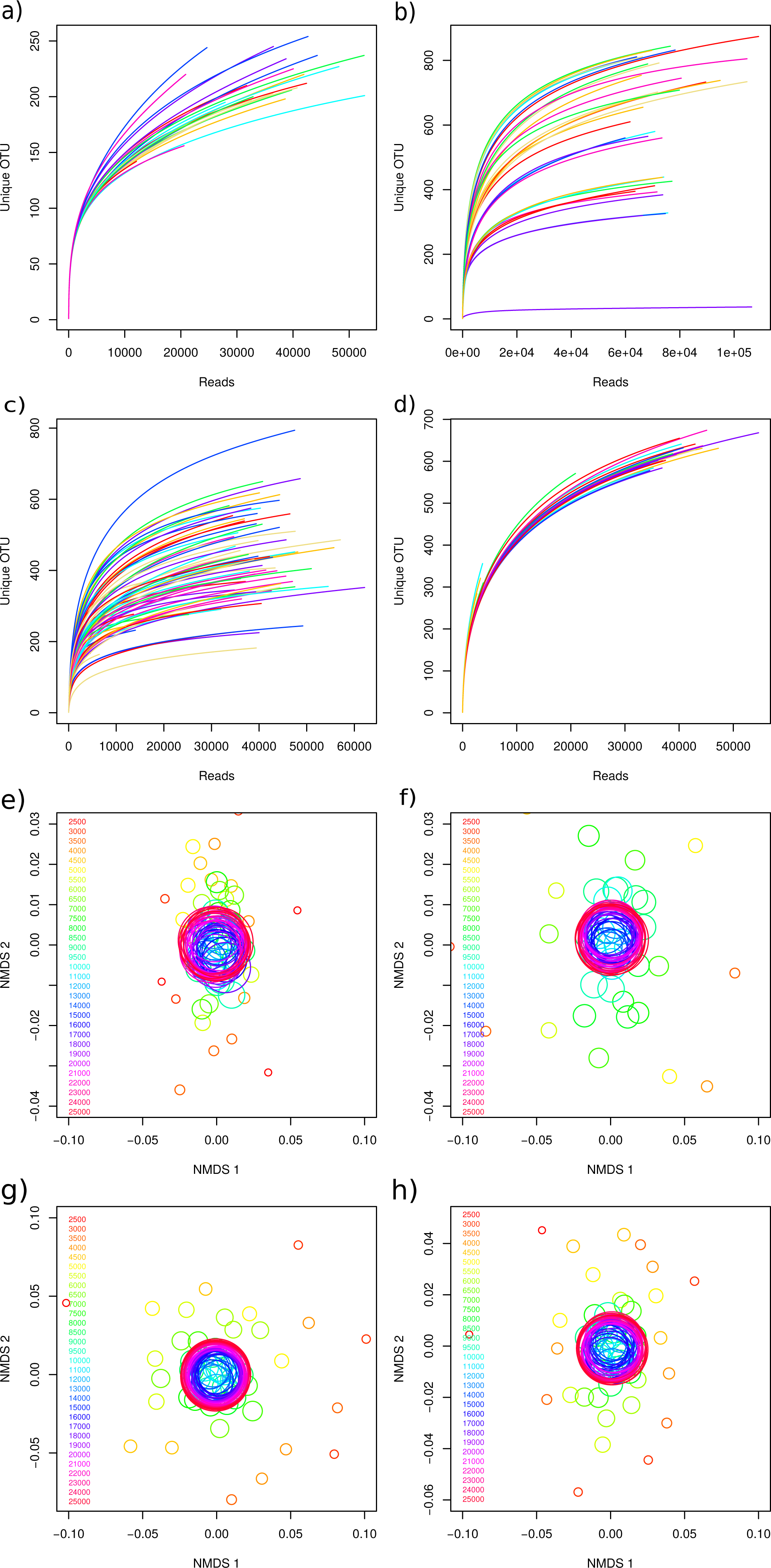
The effect of sequencing depth on data is more pronounced for assessments of richness than of beta-diversity. Richness is still growing at 50,000 reads for (a) fecal samples (b) vaginal swabs, (c) intestinal biopsies and (d) saliva samples. However, looking at beta-diversity, these same samples are stable at c. 12,000 reads (e: fecal; f: vaginal; g: intestinal; h: saliva).

However, another way to think about sequencing depth is whether enough gene tags have been read to allow the comparison of two related populations. To investigate this other point of view, the samples used in figure 6 a-d were pooled into one artificial pool per sample type. This artificial pool thus has higher richness than any individual sample, and would require a deeper sequencing effort to be understood. We then simulated samples of different sizes from these pooled samples and found that 10-12,000 high quality reads are enough to give a stable picture of these communities **(fig 6e-h)**, suggesting that the remaining OTU discovered are in very low abundance. Given the error margins involved in pooling and sequencing, aiming for 20,000 reads per sample is likely to be a safe strategy.

### OTU picking and taxonomy assignment

While 97% average-linkage clustering is a *de facto* standard of the field, OTU picking is probably one of the most controversial topics in microbiomics [28–32]. To identify a suitable approach for the host-associated environments in this study, OTU were clustered at different cutoffs or corrected by Unoise [23]. Error correction can yield a more accurate number of OTU for the mock, but a suitable cutoff for removing rare sequences is crucial **(fig 7a, b)**. As a rule of thumb, we recommend setting this cutoff equal to the number of samples, but this parameter might require fine-tuning to the microbial community of interest.

**Figure 7.**
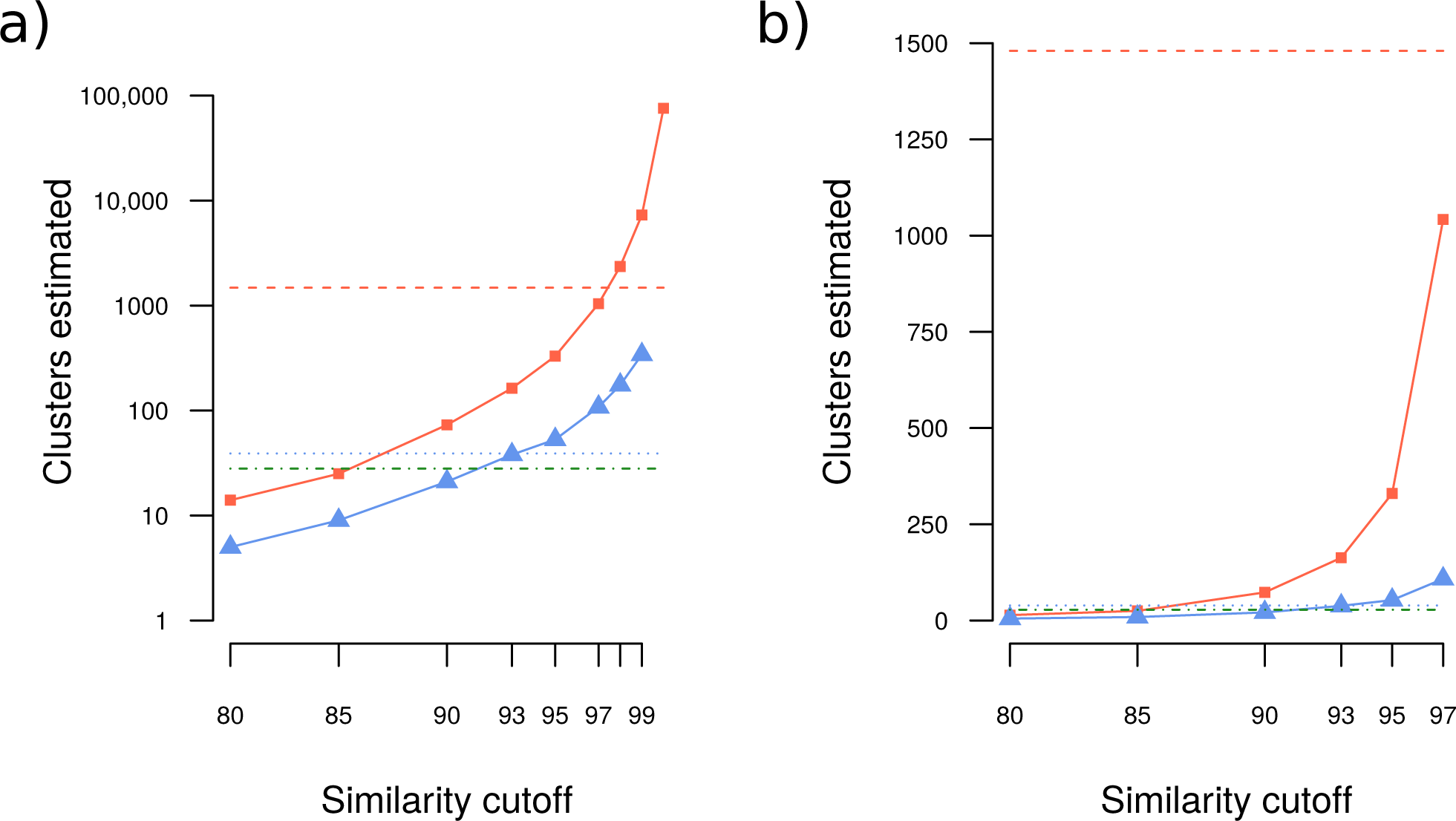
Estimated number of OTU from mock samples based on different cluster similarity cutoffs or error correction. (a) The x-axis represents different levels of clustering, from 80% to 100%. The y-axis is the number of OTU observed (log 10 scale). The continuous lines and dots are clustered at various similarity cutoffs and discard singletons only (orange line, squares) or removing OTU whose total abundance was less than the number of samples (in this case, 72; blue line, triangles). The orange dashed line corresponds to error correction with Usearch and removal of singletons, while the blue dotted line corresponds to removal of all OTU with abundance less than 72. The green line represent the expected number of OTU in these samples. (b) Same as (a), but with a linear y-axis. Only clustering identities between 80% and 97% are shown.

After OTU picking, the next logical step is to assign taxonomic annotation to each centroid sequence. Taxonomy was assigned using either the RDP classifier with its native database [25], SINA with the ARB-SILVA database [24] or an adaptation of the mapping strategy developed by Hu and coworkers [26], also using the ARB-SILVA database (see methods for details). Each of these three approaches was assessed in three ways: *in silico* simulated samples, sequencing data from the Zymo mock community (see methods) and sequencing data from faecal samples and vaginal swabs. While in the first two cases the goal is to replicate as closely as possible the underlying community structure, in the latter we aim to retrieve a reasonable number of OTU and classifying as many of them as possible to species-level resolution. SINA and the mapping strategy gave comparable results at the genus level, while the RDP classified most sequences to family level. The mapping strategy is the only one of the three assessed capable of assigning a species to any of the sequences, and had overall deeper classification than SINA (**fig 8).** This advantage is lost in environments with less representation in the available databases, but can be important for studies of the human microbiome.

**Figure 8.**
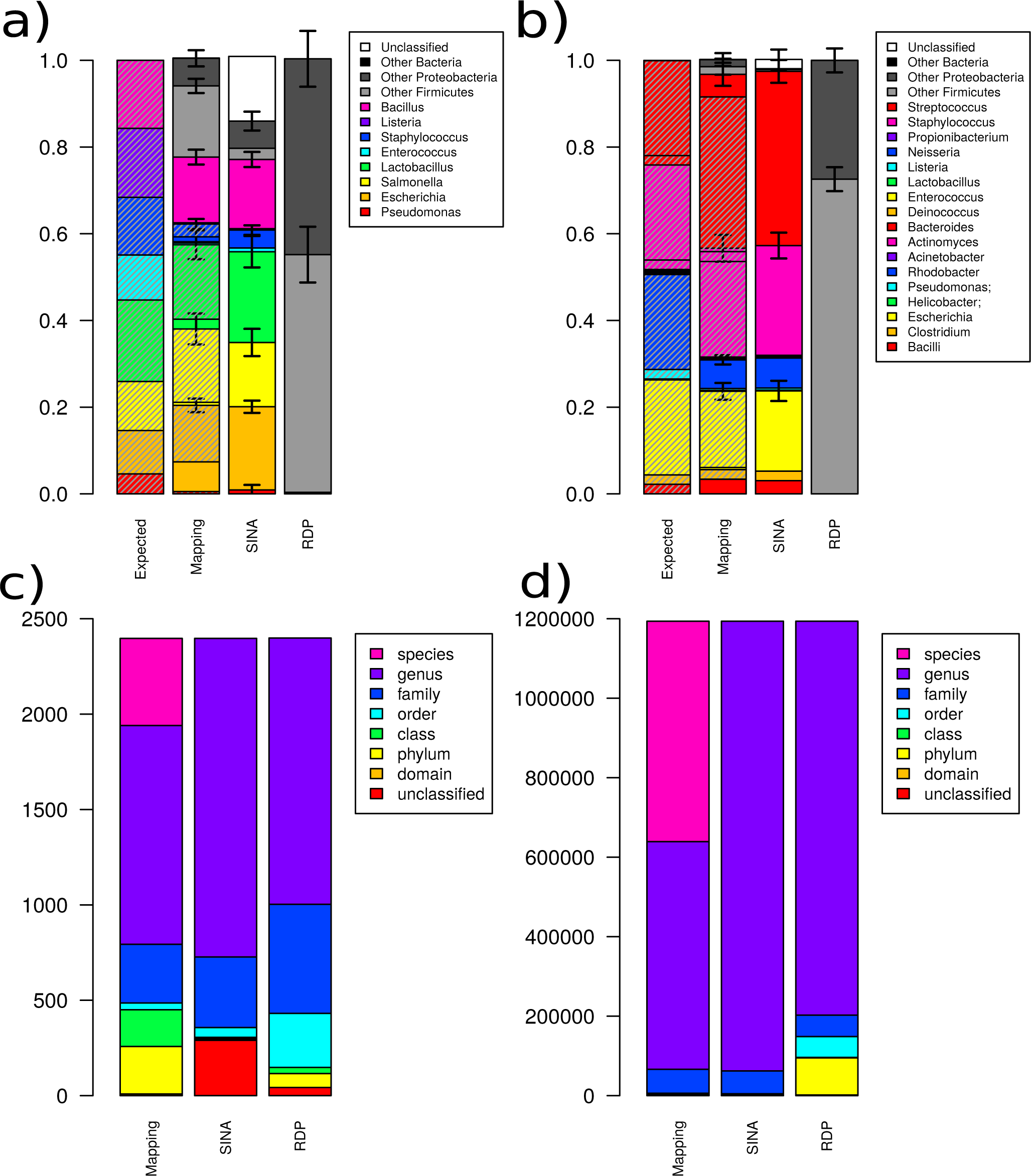
Classification depth and accuracy with different methods. (a) Several defined mock samples (“Zymo mock”) were sequenced, processed and classified with ARB-SILVA, RDP or mapped to a manually curated subset of the Silva database, as described in the methods. The mapping strategy presents the best results at the genus level, in addition to being the only approach that provides species-level information. The shaded areas over the mapping stacked bars highlight the proportion of samples classified to the correct species, as opposed to a genus-level assignment or misclassification. We notice that most of the RDP classification results were at the class level, so our choice of depicting genera and phyla is biased against this method (b) Same as (a), but with a different mock community (“BEI mock”) (c) Fecal samples, vaginal swabs and biopsies were sequenced and classified with the same methods as in (a) and (b). SINA has the highest rate of unclassified OTU. The in-house method is the only one with species-level resolution. (d) Same as (c), but scaled on the number of reads classified at each depth, rather than the number of OTU. All methods show a bias towards classifying abundant OTU in more detail, since these common clades are more likely to be found in reference databases.

## DISCUSSION

The workflow discussed here is sturdy and flexible, but like any other laboratory protocol, it will need to be adapted for the material conditions of each lab. The workflow described is optimized for a medium to high-throughput setting running a minimum of one sequencing run per week. To achieve this, infrastructure has to be scaled accordingly, including freezer space, the availability of liquid handling robots both for DNA extraction and for library preparation and a laboratory information management system (LIMS) capable of keeping track of each sample as it runs through the pipeline. Smaller labs can still adapt the pipeline to their settings, taking extra precautions to prevent contamination or material degradation during manual sample handling.

We notice that the 1-step PCR protocol, while generally superior to the 2-step, is more sensitive. This is a natural thermodynamic consequence of the length of the primers. Therefore, if the extracted DNA cannot be made sufficiently pure for the 1-step method, a 2-step can be adopted. Furthermore, specific research questions may require other primer choices (eg [5,33,34]).

Finally, while good laboratory practices such as the ones outlined here can go a long way in improving the reproducibility and applicability of human microbiome studies [35], it is still crucial to not leave behind the central lessons of epidemiology when designing cohort and case-control studies [36, 37].

## CONCLUSIONS

Here we present a flexible and scalable approach to library preparation and analysis for human microbiome studies (**fig 9)**. It can easily be applied to samples collected at home, in clinics or during field work. Its direct applicability to saliva samples, for which the method hadn’t been previously optimized, is further evidence of its robustness. It is highly reproducible and minimally biased, and can be performed manually or with the assistance of liquid handling robots. This approach can easily be extended to other tissue and sample types, providing an extensively validated suite of best-practices for microbiome studies.

**Figure 9.**
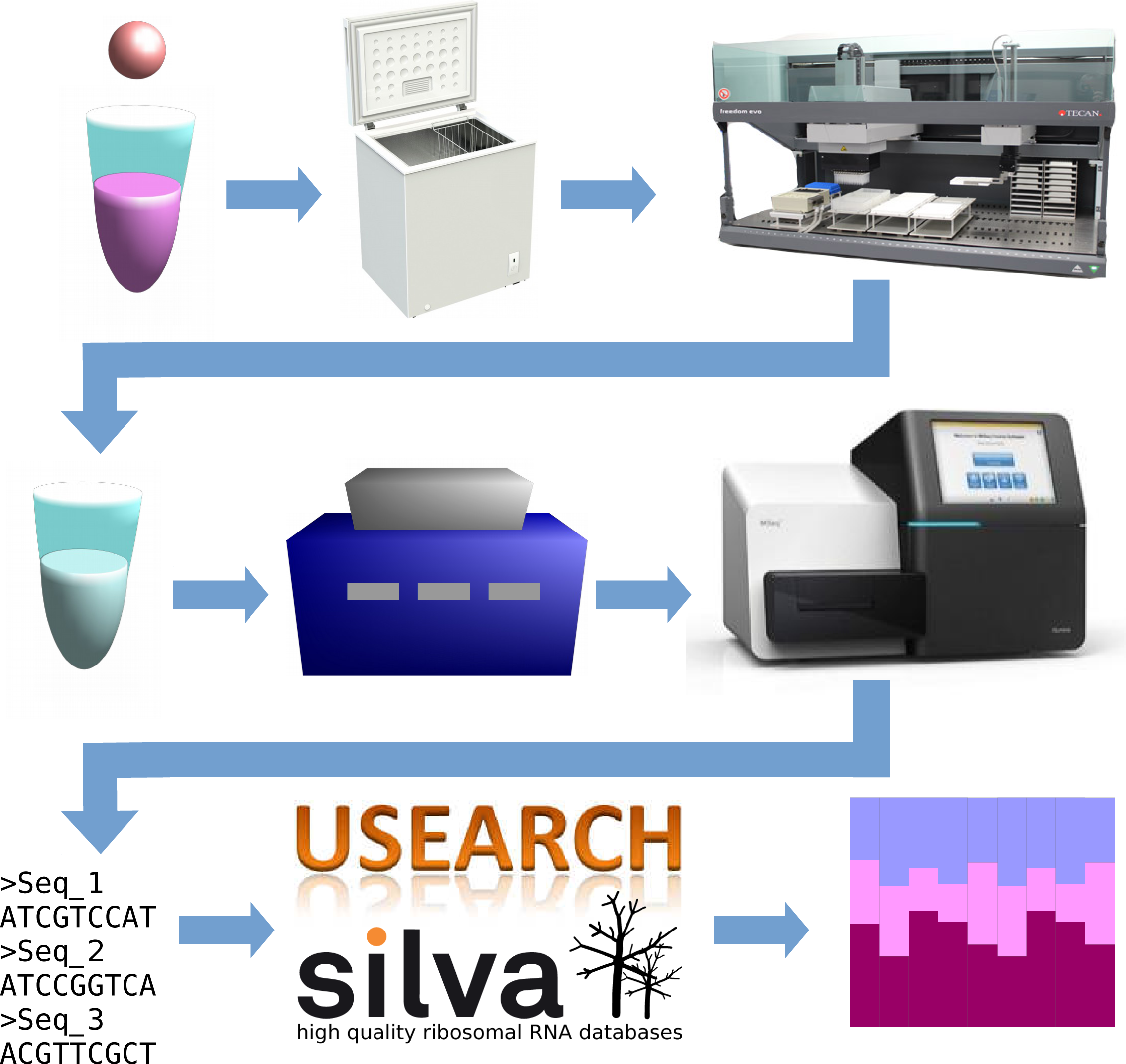
Overview of the CTMR pipeline. Any sample collected should be immediately preserved in DNA/RNA-shield. They can then be preserved for several days at room temperature or lightly frozen. An extraction pipeline comprising a sequence of physical bead-beating and chemical digestions thoroughly extracts DNA from any human-derived sample. A 1-step PCR amplification with universal primers guarantees minimal biases and produces Illumina-ready samples. A bioinformatics pipeline based on Usearch and the ARB-SILVA database then annotates this samples at species-level accuracy.

## METHODS

### Sample collection and storage

Unless otherwise stated, samples were collected, stored in DNA/RNA shield (Zymo Resarch Corp, Irvine, CA), kept at −20°C for up to 12h and then transferred to −80°C until extraction time. DNA/RNA Shield is claimed by the manufacturers to completely inactivates bacteria in under five minutes, allowing for room-temperature storage of samples. Samples thus specified were kept in RNAlater (Qiagen, Venlo, Netherlands) or Allprotect Tissue Reagent (Qiagen, Venlo, Netherlands). All biopsies used in this study were wrapped in sterile paper and fresh frozen at −80°C.

### DNA extraction

Unless otherwise stated, samples not already preserved in DNA/RNA Shield were transferred to this buffer prior to extraction with ZR-96 Genomic DNA MagPrep (Zymo Research Corp, Irvine, CA). 1 mL of DNA/RNA shield was added to 30-100 mg of fresh frozen fecal sample.

Fecal samples and biopsies in DNA/RNA shield were submitted to beating with Matrix E (MP Biomedicals, Santa Ana, CA, USA) at 1600 RPM in a 96 FastPrep shaker (MP Biomedicals, Santa Ana, CA, USA) for 1 minute (fecal samples) or 2-6 minutes (biopsies; the beating proceeds until the sample is visually homogeneous). Vaginal swabs and saliva samples were beat for 1 minute in ZR Bashing Bead lysis tubes (Zymo Research Corp, Irvine, CA, USA). Samples were spinned down at 4400 RPM for 4 minutes to remove beads from the solution. Samples were then incubated in lysozyme buffer (20 mM Tris-CL, 2 mM sodium-EDTA, lysozyme to 100 g/mL; Sigma, St. Louis, MO, USA) at 37°C for 45’ to 1h at 1000 RPM in the following proportions. For samples kept in DNA/RNA-shield, 200 µL of pre-treated sample and 20 µL of lysozyme buffer; for biopsies that had not been preserved in this way, 126 µL of sample and 14 µL of buffer were used. Faecal samples were incubated an additional 15 min at 80°C and 250 RPM. Following this, samples were spinned down for 5’ at 4400 RPM and 200 µL are transferred to a new plate, to eliminate larger particles. 10 µL of proteinase K (from the Genomic DNA MagPrep kit, Zymo Research Corp, Irvine, CA, USA; the 2x digestion buffer in the kit is not necessary if DNA/RNA-shield is used) added, and they are incubated at 55°C at 250 RPM for 30 minutes. The samples are then cleaned through several washing and magnetic bead peletting steps according to the instructions of the manufacturer (Genomic DNA MagPrep kit, Zymo Research Corp, Irvine, CA, USA).

Unless otherwise stated, samples were eluted from the magnetic beads with 70 µL of Elution Buffer (10 mM Tris-Cl, ph 8.5; Qiagen, Venlo, Netherlands). Otherwise, PCR-grade water (brand) or buffer TE (10 mM tris-Cl, 1mM EDTA, ph 8.5; Qiagen, Venlo, Netherlands) were used as described in the results section. Samples are incubated for 10 minutes at 65°C in Elution Buffer and then peletted again with a magnet. The pure DNA is now in the supernatant. The steps described in this paragraph can be performed manually, but have been automated to be performed by a FreedomEVO robot (TECAN Trading AG, Männendorf, Switzerland).

Samples extracted with the MoBio PowerMag Microbiome RNA/DNA isolation kit (Qiagen, Venlo, Netherlands) were processed according to instructions by the manufacturer. A blank negative control and a positive mock control (ZymoBIOMICS Mock Community Standard, Zymo Research Corp, Irvine, CA, USA) have been included in each extraction round. Unless otherwise stated, lysis was performed with lysozyme (Sigma-Aldrich, Saint Louis, MO); in the cases thus specified, the lysing agent used was BugLysis (Molzym GmbH & CO KG, Bremen, Germany).

### DNA amplification and sequencing

Prior to amplification samples were normalized and a total of 50 ng for samples with low eukaryotic content (fecal samples, vaginal swabs and saliva samples) or 170 ng for samples with high eukaryotic content (biopsies), have been used to amplify the V3-V4 region of the 16S rRNA gene using primer pair 341F/805R [38].

For the 1-step PCR procedure, amplification was carried out by a high fidelity proof-reading polymerase for a total of 20 cycles (fecal samples and vaginal swabs) or 25 cycles (biopsies). For amplification of the sequencing libraries, forward primer 5’-CAAGCAGAAGACGGCATACGAGAT-N8-GTCTCGTGGGCTCGGAGATGTGTATAAGAGACAGGACTACHVGGGTATCTAATCC-3’ and reverse primer 5′AATGATACGGCGACCACCGAGATC-N8-TCGTCGGCAGCGTCAGATGTGTATAAGAGACAGCCTACGGGNGGCWGCAG-3’, where N8 represents an identifying 8-mer (barcode) and the last 21 and 19 bases in each construct are the sequence specific forward and reverse primers, respectively, were used.

For the 2-step procedure, the initial amplification was carried out for 20 cycles (fecal samples and vaginal swabs) or 25 cycles (biopsies), with the forward primer construct 5’-TCGTCGGCAGCGTCAGATGTGTATAAGAGACAG**CCTACGGGNGGCWGCAG**-3’, comprising the 341F universal bacterial primer sequence (in bold); and a Illumina specific adapter overhang sequence and the reverse primer construct and the reverse primer construct 5’-GTCTCGTGGGCTCGGAGATGTGTATAAGAGACAGGAC**TACHVGGGTATCTAATCC**-3’, comprising the universal bacterial primer 805R (in bold) and a Illumina specific adapter overhang sequence. After cleaning, a 5 µL aliquot of each of the 2-step samples was submitted to an Indexing reaction using the Nextera XT Index Kits v2 (Illumina Inc, San Diego, CA, USA), in a 13 cycle PCR.

Purification of PCR products was carried out using Agencourt Ampure XP Beads (Beckman Coulter AB, Stockholm, Sweden) on a KingFisher Flex System. Finished libraries were quantified using Invitrogen’s QuantIt fluorometric assay (Thermo Fischer Scientific, Waltham, Massachusetts, USA). Samples were then pooled to equimolar amounts and sequenced in parallel to whole bacterial genomes in a MiSeq instrument (Illumina Inc, San Diego, CA, USA). All controls from the extraction phase, as well as a negative (blank) PCR control have been submitted to PCR and consequently sequenced with the respective samples.

### Sequence correction and taxonomic assignment

Each sequencing run was processed independently, together with all other 16S projects in the same run. Cutadapt [39] was used to eliminate all sequences not containing the amplification primers, remove the primer sequences, all bases with a Phred score below 15 and all reads with less than 120 bp left after trimming. The resulting reads were merged with Usearch v.9.0.2132[40] and reads failing to merge or producing merging products shorter than 380 bp, longer than 520 bp or with more than three expected errors were discarded. All unique full-length samples occurring at a frequency higher than 10^−6^ in the dataset were submitted to Unoise [23]. Alternatively, Usearch was used to cluster them at 97% or 99% identity, as described in the Results section. All the merged reads were then mapped back to the accepted centroids and assigned to the OTU with the highest identity, at a minimum of 98%. In case of equally high identity matches, the most abundant centroid is selected. Taxonomy was assigned using either SINA v.1.2.13 [24], the RDP online classifier [25] or an adaptation of the mapping method described by Hu et al [26]. In the latter case, reads were mapped at a minimum of 89% identity to the SILVA 129 database. The taxonomic annotation of the database was manually curated to remove uninformative species-level information (e.g. “soil metagenome”) as well as obvious misclassifications (such as eukaryotic species annotated as bacteria). For the classification levels of strain, species, genus, class and phylum, identity cutoffs were set at 100%, 99%, 97%, 95% and 90%, respectively. The top hit above each identity threshold was identified and all hits with scores at least 95% of it and still above the cut-off were selected. The last common ancestor of these taxa is then selected and truncated at the appropriate taxonomic level. The most complete classification for each OTU is finally selected. Since the curated database does not include strain information, this classification level is in practice not done, but is kept to assure a preference for exact matches over 99% identity matches. Scripts for performing this analysis are available at https://github.com/ctmrbio/Amplicon_workflows. For statistical analyses, all samples with fewer than 2000 high-quality reads were discarded, as were OTU present in fewer counts than the number of samples under analysis.

## DECLARATIONS

### Ethics approval and consent to participate

All samples used in this work were collected from studies approved by the local ethics committees in Uppsala, Sweden (Dnr. 2008/395 for gut biopsies on filter paper, Dnr 2016/517 for vaginal and fecal samples) and Stockholm, Sweden (Dnr. 394/01 for gut biopsies in the main text and Dn.r 2015/2138-31/2 for gut biopsies in the supplementary figure).

### Consent for publication

NA

### Availability of data and materials

The datasets discussed in this work are available at the NCBI short read archive under the identifiers XXXXXXXXXX. The code used for the early data analysis and a step-by-step tutorial is available at https://github.com/ctmrbio/Amplicon_workflows, version April 4th. Improvements to this workflow will be added to the same repository. The R code used to generate the figures is available as Supplementary File 1 and can also be provided by the first author in plain text or jupyter notebook formats, as needed.

### Author’s Contributions

AALP, MS and JD conducted the experiments. AALP, MS, MCH and LWH participated in the planning and analysis of the experiments. LWH performed all bioinformatic and statistical analyses. LWH, AALP and MS wrote the paper. All authors read and approved the final manuscript.

## Ackwnowledgements

The study has been supported by the Center for Translational Microbiome Research (CTMR), a collaboration between Karolinska Institutet, Science for Life Laboratory and Ferring Pharmaceuticals. We further acknowledge Niklas Nyström for providing us with biopsies of the development of the method.

---

**Supplementary figure 1.**
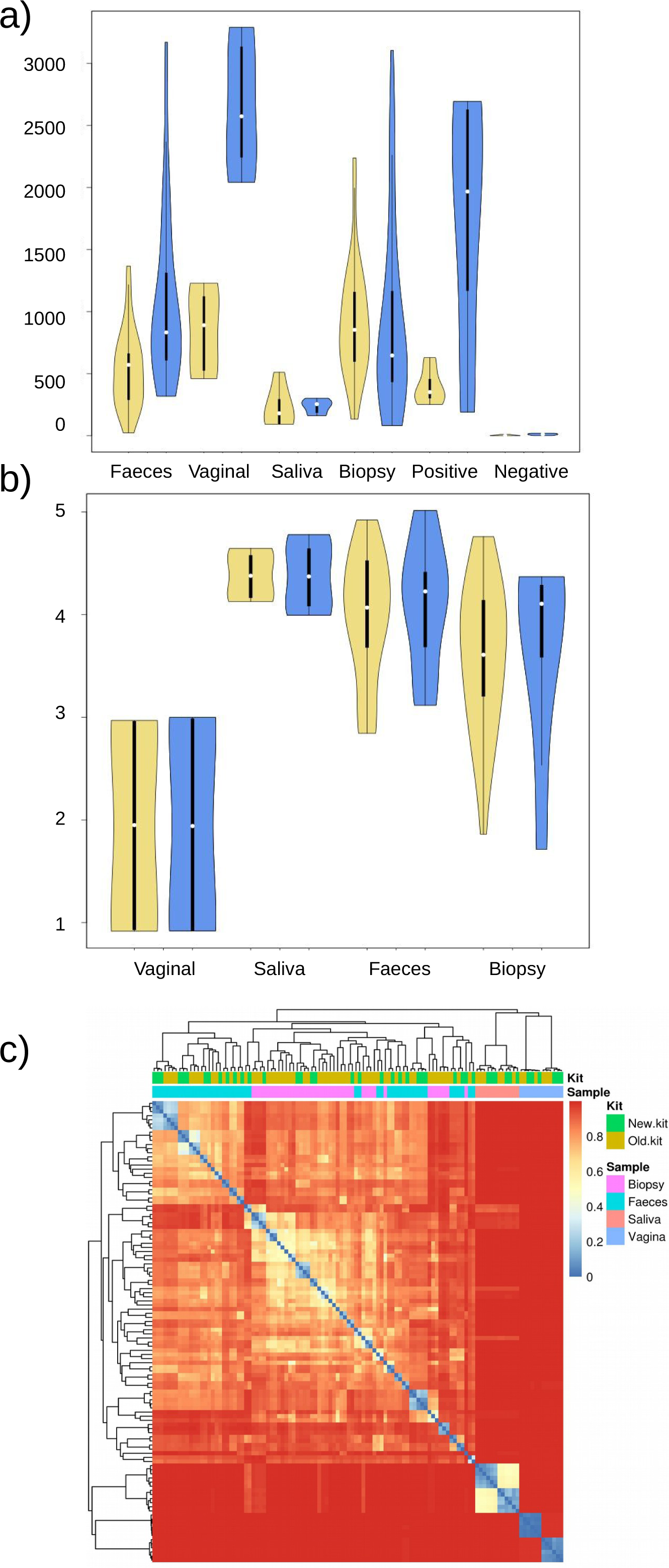
In plots (a) and (b), yellow corresponds to the ZR-96 Genomic DNA MagPrep kit and blue to the Quick-DNA MagBead Plus. Each sample type is marked in the x-axis. (a) DNA yield in ng. (b) Shannon’s alpha-diversity. (c) Heatmap of Bray-Curtis divergence between samples. The coloured lines above the heatmap indicate the kit used (Old.kit = ZR-96 Genomic DNA MagPrep; New.kit = Quick-DNA MagBead Plus) and the sample type. Biopsy and faecal samples don’t entirely segregate, but the saliva and vaginal samples form their distinct clusters. Within each cluster there is a minor effect of the kit used, so different kits should not be mixed within the same analysis.

